# cryoTIGER: Deep-Learning Based Tilt Interpolation Generator for Enhanced Reconstruction in Cryo Electron Tomography

**DOI:** 10.1101/2024.12.17.628939

**Authors:** Tomáš Majtner, Jan Philipp Kreysing, Maarten W. Tuijtel, Sergio Cruz-León, Jiasui Liu, Gerhard Hummer, Martin Beck, Beata Turoňová

**Affiliations:** Department of Molecular Sociology, Max Planck Institute of Biophysics, Frankfurt am Main, Germany; IMPRS on Cellular Biophysics, Frankfurt am Main, Germany; Department of Theoretical Biophysics, Max Planck Institute of Biophysics, Frankfurt am Main, Germany; Institute of Biophysics, Goethe University Frankfurt, Frankfurt am Main, Germany; Institute of Biochemistry, Goethe University Frankfurt, Frankfurt am Main, Germany

## Abstract

Cryo-electron tomography enables the visualization of macromolecular complexes within native cellular environments, but is limited by incomplete angular sampling and the maximal electron dose that biological specimen can be exposed to. Here, we developed cryoTIGER (Tilt Interpolation Generator for Enhanced Reconstruction), a computational workflow leveraging a deep learning-based frame interpolation to generate intermediate tilt images. By interpolating between tilt series projections, cryoTIGER improves angular sampling, leading to enhanced 3D reconstructions, more refined particle localization, and improved segmentation of cellular structures. We evaluated our interpolation workflow on diverse datasets and compared its performance against non-interpolated data. Our results demonstrate that deep learning-based interpolation improves image quality and structural recovery. The presented cryoTIGER framework offers a computational alternative to denser sampling during tilt series acquisition, paving the way for enhanced cryo-ET workflows and advancing structural biology research.

## 1 Introduction

Cryo-electron tomography (cryo-ET) has revolutionized our ability to visualize macromolecular complexes within their native cellular environments. At the heart of cryo-ET is the tilt series acquisition process, where a biological specimen is imaged at incremental tilt angles to generate a series of two-dimensional (2D) projections. These projections are then computationally reconstructed into a three-dimensional (3D) volume called tomogram, providing insights into the structural organization of cellular components^1^.

The acquisition setup requires optimizing the interplay of multiple parameters at once to obtain tilt series with desired qualities. The most crucial parameters are the total electron dose imposed on the sample, the tilt range, and the tilt increment. Since biological samples are highly sensitive to radiation damage, excessive electron dose will degrade the sample, compromising the integrity of the structural information^2, 3^. The tilt range determines the effective thickness of the sample during tilting and, more importantly, the extent of the missing wedge (i.e., the angular space with missing signal)^4–6^.

Finally, the tilt increment, the angular step between successive projections, directly influences the completeness of angular sampling and thus the completeness of the 3D reconstruction. The relationship between the angular sampling and the resolution beyond which the signal content becomes incomplete is described by the Crowther criterion^7^. Smaller increments provide more complete angular sampling, thereby enhancing the contrast and visibility of smaller features^4, 8, 9^. To maintain a reasonable tilt range, one has to either increase the total electron dose or decrease the dose per tilt, both of which complicate the subsequent processing by lowering the signal-to-noise ratio (SNR) for each image^10^. Conversely, larger increments allow for a higher electron dose per tilt but lead to poorer angular sampling and stronger artifacts in the tomograms^5, 11^.

In standard practice, tilt series are typically acquired at increments of two or three degrees with tilt range *±* 60 degrees^12^. This setup has proven itself well-suited for obtaining high-resolution structures using the subtomogram averaging (STA) workflow in which multiple instances of the same complex are localized within tomograms and then aligned and averaged together^10^. The aligning and averaging of randomly oriented particles effectively extends the angular sampling and thus reduces missing wedge in the obtained structure.

A crucial step of STA is reliable particle localization, which remains challenging especially for smaller complexes. The most common localization methods are template matching^13–15^, deep-learning (DL) based approaches^16–19^, and surface-based localization that is used for pleomorphic assemblies^20–23^. While the negative impact of the missing wedge on the depiction of features that are elongated perpendicularly to the beam direction has been well described^24, 25^, the extent to which the incomplete angular sampling between the tilts negatively influences those methods remains understudied. Consequently, most of the research is focused on filling the missing wedge^26–28^ while the incomplete angular sampling between tilts has not been systematically explored.

When looking at angular sampling from the perspective of computer vision, one wants to synthesize intermediate images between a pair of input tilts with a certain motion. In general, it is possible to address this with traditional methods based on linear or tricubic interpolation^29^ or more advanced DL-based image interpolation techniques^30^. The latter leverage the power of convolutional neural networks (CNNs)^31^, recurrent neural networks (RNNs)^32^, and generative adversarial networks (GANs)^33^ to learn representations of image content and spatial relationships. DL models are trained on large datasets, allowing them to capture a wide range of textures and motions displayed in the field of view. To the best of our knowledge, none of these interpolation methods have been applied so far to generate additional images within cryo-ET tilt series.

Here we present cryoTIGER: Tilt Interpolation Generator for Enhanced Reconstruction for cryo-ET, which computationally reduces the angular spacing by interpolating between the neighboring images within the tilt series. We adapted a DL-based frame interpolation algorithm called FILM^34^ for the cryo-ET workflow and trained our own models on multiple datasets, providing sufficient diversity in acquisition parameters and cellular content. We evaluated a pre-trained and our own models and compared them to linear interpolation as well as to non-interpolated data. The results showed that in comparison to non-interpolated data, our DL-based interpolation yielded better outcomes in most use cases. Our study underlines the importance of more complete angular sampling between the tilts and provides a computational solution which reduces the need to physically acquire datasets with exceedingly dense angular sampling.

## 2 Results

### 2.1 Adaptation to the cryo-ET workflow

To interpolate artificially generated tilt-images in between experimentally acquired tilts, we choose Frame Interpolation for Large Motion (FILM)^34^. For technical details we refer the reader to the original work. Briefly, FILM employs a multi-scale feature extractor^35^ that shares weights across the scales and presents a ‘scale-agnostic’ bidirectional motion estimation module. This approach relies on the notion that large motion at finer scales should be similar to small motion at coarser scales, thus increasing the number of pixels available for large motion supervision. Similar to how the module ensures consistent motion representation across varying scales, this principle can be applied to tilted images, where key features must remain recognizable despite geometric distortions. By maintaining adaptability and efficiency across transformations, the method aligns well with the challenges of our setup, ensuring robust performance under varying tilt angles. In order to accommodate the FILM framework for cryo-ET data, we made multiple adjustments, as shown in Fig. 1.

**Figure 1.**
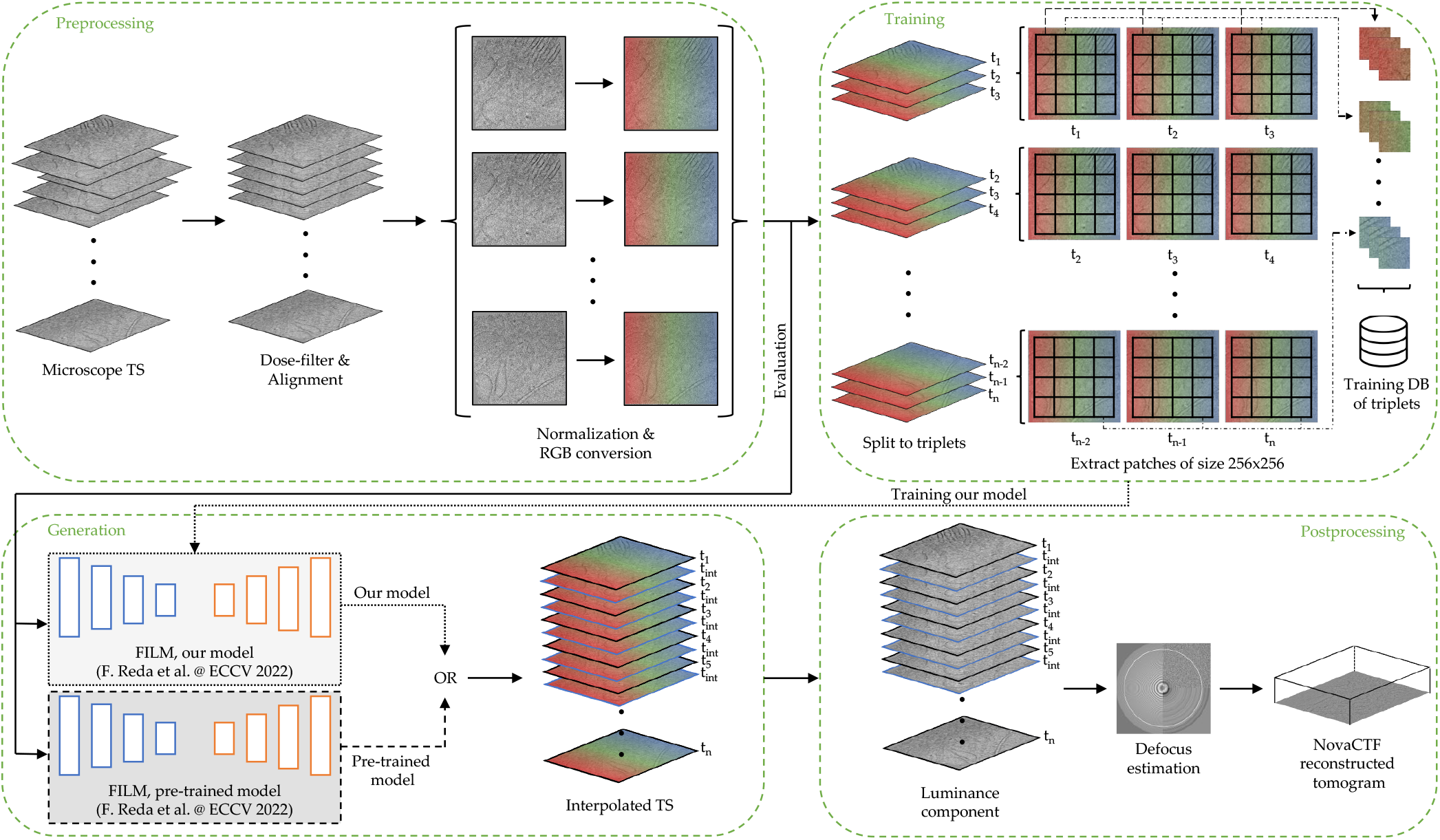
Pipeline of cryoTIGER depicting incorporation of a deep learning frame interpolation model into cryo-ET reconstructions. Individual steps are described in the main text.

Our tilt-series processing pipeline consists of *Preprocessing, Training, Generation*, and *Postprocessing* steps. In *Preprocessing*, raw microscope tilt series images undergo the basic operations of dose filtering, alignment, normalization, and conversion from the gray-scale images to colored images with red, green, and blue (RGB) channels in order to make them compatible with the FILM framework. During the *Training* step, we utilised data from multiple cryo-ET datasets, see Table 1. The training data were collected with different pixel sizes and tilt increments to increase the robustness of our DL model. The majority of these tilt images were acquired with more commonly used increment of two or three degrees but we included a one degree tilt increment dataset as well, in order to have items with smaller motion in the training dataset. Supplementary Table 1 provides a list of models that we considered and trained for this work. When we trained the model on larger tilt increments only, the performance slightly worsened (see Supplementary Fig. 1). From now on we refer to *our model* as the model trained on 317,312 triplets from 375 tilt series that is highlighted in Supplementary Fig. 2b.

**Table 1.** Table summarizing the pixel size, tilt increment, and number of tilt series that were used to train our model.

| Sample | Pixel Size | Tilt Increment | # of TS |
| --- | --- | --- | --- |
| *D. discoideum* | 1.971 | 1 | 49 |
| *D. discoideum* | 1.971 | 2 | 77 |
| *D. discoideum* | 1.971 | 3 | 159 |
| Human T cells | 1.971 | 2 | 32 |
| Human kidney cells (HEK Flp-In T-REx) | 1.971 | 2 | 10 |
| Human kidney cells (HEK-293) | 1.188 | 2 | 18 |
| Human skin fibroblasts | 2.414 | 2 | 30 |
| **SUM** |  |  | **375** |

The FILM framework proposed a unified architecture for image interpolation, which is trainable from regular image triplets alone^34^. In our setup, a triplet refers to a set of three consecutive tilt images, where the two external ones are used to interpolate a tilt image between them and the middle tilt image is used as the ground truth image for comparison. Therefore, in order to train the network with our inputs, we first split tilt images into triplets. Because the input size of triplets for training using FILM framework is 256 × 256, we further divided each tilt into patches of this size and stored them in a training database.

In *Generation* step, we tested both a pre-trained FILM model and a model trained using our data to interpolate additional tilts in between the ones acquired physically. During the *Postprocessing*, the luminance component representing overall brightness of an image is extracted (see Methods 4.1 for details), followed by defocus estimation using Gctf^36^, and tomographic reconstruction using novaCTF^37^, which performs correction of contrast transfer function (CTF) in 3D. Note that since we use aligned tilt series for interpolation the generated images do not require any alignment prior the reconstruction.

### 2.2 Analysis of 2D interpolated tilt images

We first analyzed the generated data in 2D by comparing their quality to the available ground truth (GT) images. We used *Dictyostelium discoideum* tilt series acquired with a one-degree tilt increment (see Table 1) and split them into even and odd tilts. Odd tilts, starting from index one, were used as input to generate interpolated tilts, while even tilts, starting from index two, served as GT data.

We investigated three different interpolation approaches and compared them to the GT. The first approach is a linear interpolation model, in which we calculate the pixel-wise average between each pair of adjacent tilt images. This method serves as a simple baseline model without any deep learning component. We also considered cubic and tricubic interpolation models but their performance was worse compared to the linear interpolation model (see Supplementary Fig. 3). The remaining two approaches utilize the FILM framework to generate interpolated samples, where we employed either a model pre-trained on real-world samples from the Vimeo-90K dataset^38^ or our model as described above.

Fig. 2 compares interpolation methods against GT data. We first examined visual accuracy where all tested methods generated realistic outputs (panel A). Our model produced slightly more blurred tilts but with higher contrast compared to the other two methods. Regarding CTF estimation, the linear interpolated image showed most resemblance to the GT, while the pre-trained model and our model exhibit lower fitting accuracy (panel B). Our model exhibits more artifacts in Fourier space which is caused by noise present in the microscopic images used for the training (see Supplementary Fig. 4 and Fig. 5). Generally, the defocus values of the interpolated images are closer that of the GT (panel C), with the median defocus difference being around 50 nm, while for the pre-trained and our model the median is 100 nm and 160 nm, respectively.

**Figure 2.**
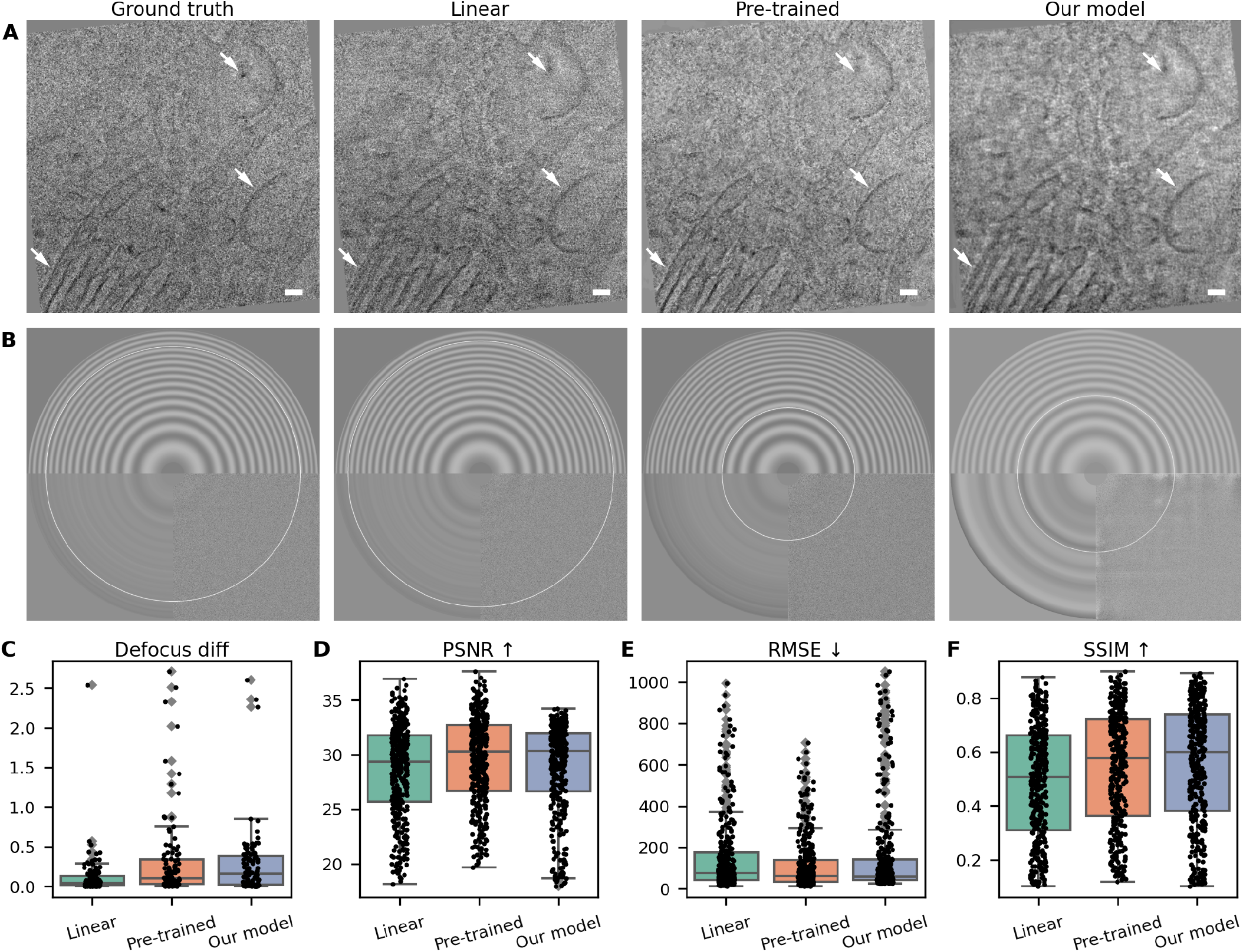
Comparison of 2D tilt series images generated using different interpolation methods. **(A)** Visual difference between features in GT image vs. interpolated images. The white arrows indicate the structural features that exhibit more pronounced differences between the tested approaches. **(B)** Defocus estimation is indicated in the Gctf diagnostic output. The white circle denotes the resolution up to which the CTF was reliably fitted. **(C)** Difference in defocus values in µm. **(D)** Peak Signal-to-Noise Ratio (PSNR), where higher values are better. PSNR is measured in decibels. **(E)**: Root Mean Square Error (RMSE), with lower values indicating better performance. **(F)** Structural Similarity Index (SSIM), with higher values indicating the images are perceived visually as more close to the original image. The scale bar is 50 nm.

For assessing visual similarity quantitatively, we employed three metrics. Firstly, the Peak Signal-to-Noise Ratio (PSNR), which measures the ratio of maximum signal power to noise power. Secondly, the Root Mean Square Error (RMSE), which calculates the square root of the average squared differences between predicted and actual values, weighting larger errors more heavily. Lastly, the Structural Similarity Index (SSIM), which evaluates perceived image quality by comparing luminance, contrast, and structural information, with values ranging from -1 (inverse structure) to 1 (identical images). See Methods 4.2 for mathematical definitions of all three metrics and description of their units. Fig. 2D-F shows comparisons between GT tilts and interpolated tilts using PSNR, RMSE, and SSIM, respectively. For PSNR, DL-based models achieve values closer to GT compared to linear interpolation, which suggests that the generated tilt series are more similar to the GT tilt images. The RMSE analysis indicates that our model exhibits a higher prevalence of outliers compared to the pre-trained model. However, the SSIM evaluation demonstrates a slight performance improvement in structural similarity for our model in comparison to the pre-trained one. Although none of the presented interpolation methods was clearly superior, the results overall highlight that we can faithfully simulate microscope tilt images using any of them.

### 2.3 Template matching on tilt-series with interpolated tilts

To evaluate the performance of our approach in generating realistic intermediate tilt images, we integrated generated images into our reconstruction pipeline (Fig.1). To quantify the impact of the interpolation for particle identification, we used high-confidence 3D template matching (TM)^15^ as implemented in GAPSTOP^TM14, 39^. We used a subset of four tilt series from the dataset EMPIAR-12454^40^ and applied TM to localize nuclear ring subunits of the nuclear pore complex (NPC NR). As a ground truth, we used the manually curated list of particle positions provided by the authors^40^. We further used a subset of four tilt series from dataset EMPIAR-11899^41^ to demonstrate the performance on the 80S ribosome using again the particle positions provided by authors of the original study. We computed the F1 score, precision, recall, and the area under the precision-recall curves (PR-AUC) to assess the performance of TM using different interpolation types (see Methods 4.3 for details).

The evaluation was conducted under two experimental scenarios. In the first scenario, we started with a tilt series acquired with a tilt increment of two degrees. We then removed every second tilt, resulting in a reduced tilt series with a tilt increment of four degrees, which served as a baseline. Subsequently, we interpolated one tilt between each pair of remaining tilts, thereby constructing a tilt series with an increment of two degrees. We also evaluated interpolation of more than one tilt but the performance typically decreased with additional interpolated tilts (see Supplementary Figure 6).

This setup allowed us to compare three distinct conditions: (1) the baseline tilt series with missing tilts after the removal step, (2) the interpolated tilt series where missing tilts were replaced with interpolated samples to restore the original number of tilts, and (3) the GT tilt series acquired directly from the microscope, which contained all the original tilts prior to the artificial removal step (see Supplementary Fig. 7 for power spectra depiction for all three cases). By comparing these conditions, we assessed the extent to which the interpolation improved tomogram properties and subsequent downstream analyses relative to the baseline without interpolation. Additionally, we compared the results to the GT reconstruction.

The results for restoring removed tilts are presented in Fig. 3A for NPC NR and in Fig. 3B for 80S ribosome. F1 scores and precision-recall graphs are illustrated on a representative tomogram, while the PR-AUC values were extracted for four tomograms from each dataset, offering a broader and more representative overview of the model performance.

**Figure 3.**
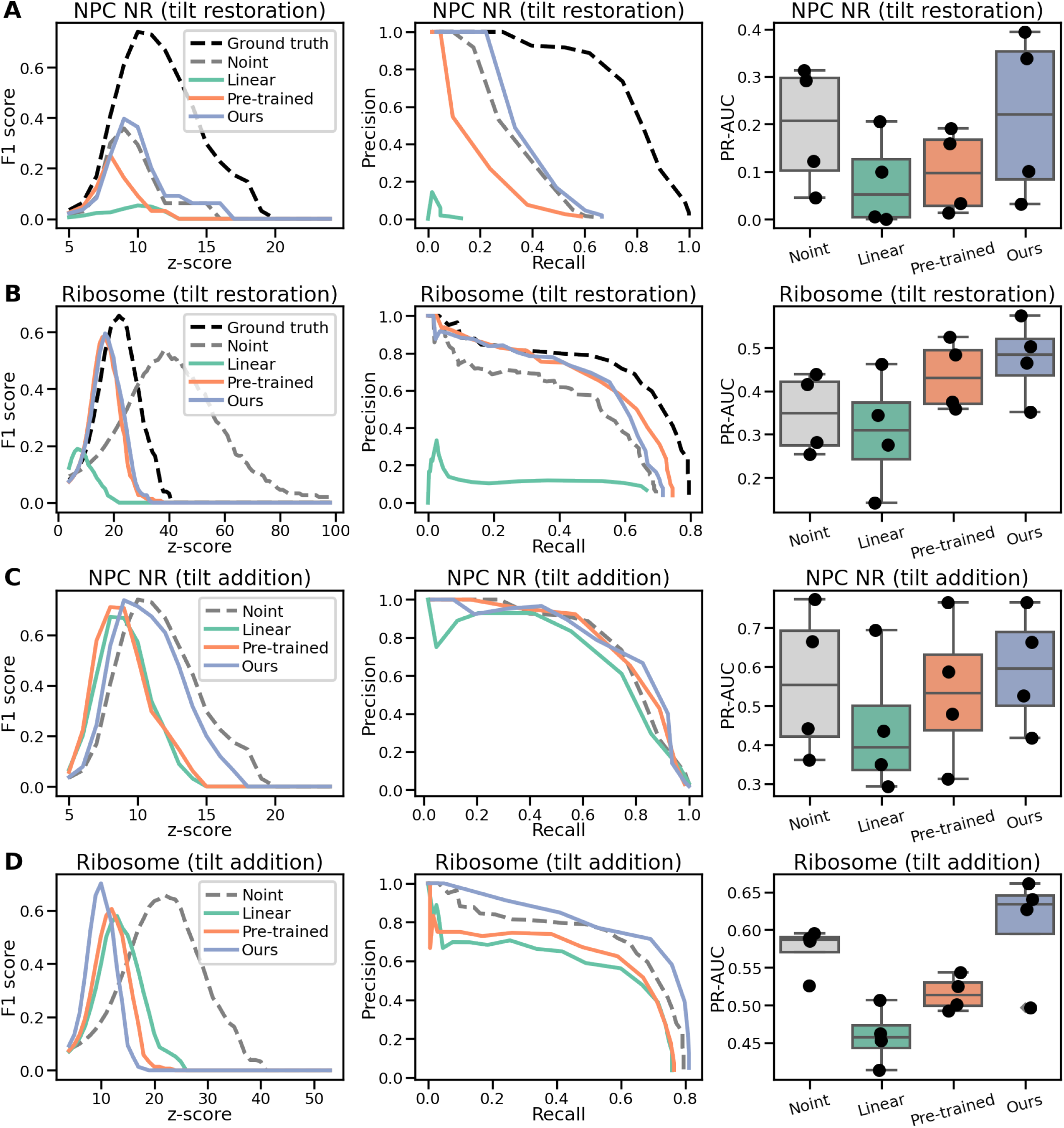
Comparison of TM results for restoring and adding tilt images using interpolation. **(A)** Results for restoring removed tilt images in the NPC NR dataset. F1 scores (left) and precision-recall curves (middle) are shown for a single representative tomogram. The area under the precision-recall curve (PR-AUC) was calculated across four tomograms (right). **(B)** Same as (A), but for tomograms containing 80S ribosomes. **(C)** Results for the addition of new interpolated tilt images in the NPC NR dataset. **(D)** Same as (C), but for tomograms containing 80S ribosomes.

It is important to note that the removal of tilts introduced a dose distribution that is to some degree artificial. In case of acquiring data with 4 degrees tilt increment one would either keep lower total dose, retaining more high-resolution content, or one would use more dose per image to achieve better contrast. In our case, the total dose is not reduced and the retained tilt images have low SNR, thereby making the interpolation task more challenging. Nevertheless, we consider this approach to be the closest approximation to a direct comparison between reconstructions with and without interpolation data.

In the second scenario, additional interpolated tilts were added to an experimental dataset, and the same set of comparisons was performed. Note that in this case the non-interpolated data contains fewer images in the tilt-series because we interpolated tilts that were not acquired. However, it will account for a more realistic experimental electron dose. The results are presented in Fig. 3C,D in the same order as for the first scenario.

Linear interpolation performed the worst in all tested cases. Notably, in the tilt restoration test, it significantly degraded the signal content, even compared to the non-interpolated data. Similar results were observed with the pre-trained model, although the differences to the non-interpolated data were less pronounced. It outperformed the non-interpolated data in the case of tilt image restoration for ribosome data. Our model outperformed the others in tilt image restoration and consistently performed better than the non-interpolated data, although the improvements were marginal in the case of NPC NR. Overall, our model demonstrated superior particle picking compared to the other tested methods and the non-interpolated variant.

### 2.4 Improving the 3D structure of nucleosomes using cryoTIGER

Encouraged by these results, we next applied cryoTIGER to enhance the localization of nucleosomes. Note that the full analysis of nucleosomes is main focus of a separate study^42^ (submitted) and here we present only improvements introduced by our framework. Nucleosomes have a molecular weight of approximately 250 kDa which, together with the crowded environment present in the nucleus and the lack of visual validation of their position, makes them an especially challenging target to identify in tomographic data. We analyzed a dataset of 14 tomograms acquired from the nuclear periphery of T cells with a tilt step of two degrees. We employed cryoTIGER to interpolate additional tilt images in between the experimental ones, resulting in tilt series with a tilt step of one degree using the pre-trained model.

The use of the interpolated tilt series lead to improved contrast in tomograms in comparison to those created from non-interpolated tilt series. As a result, individual features became more visible even without additional denoising (see Fig. 4A). This is also confirmed by Standard Deviation Contrast (non-interpolated data 0.0109 and interpolated data 0.0217) that quantifies how much the pixel values deviate from the average intensity across the entire image and by Gradient-Based Contrast (0.0034 and 0.0061, respectively) that measures the intensity variation by looking at how pixel values change between neighboring pixels (see Methods 4.4 for details).

**Figure 4.**
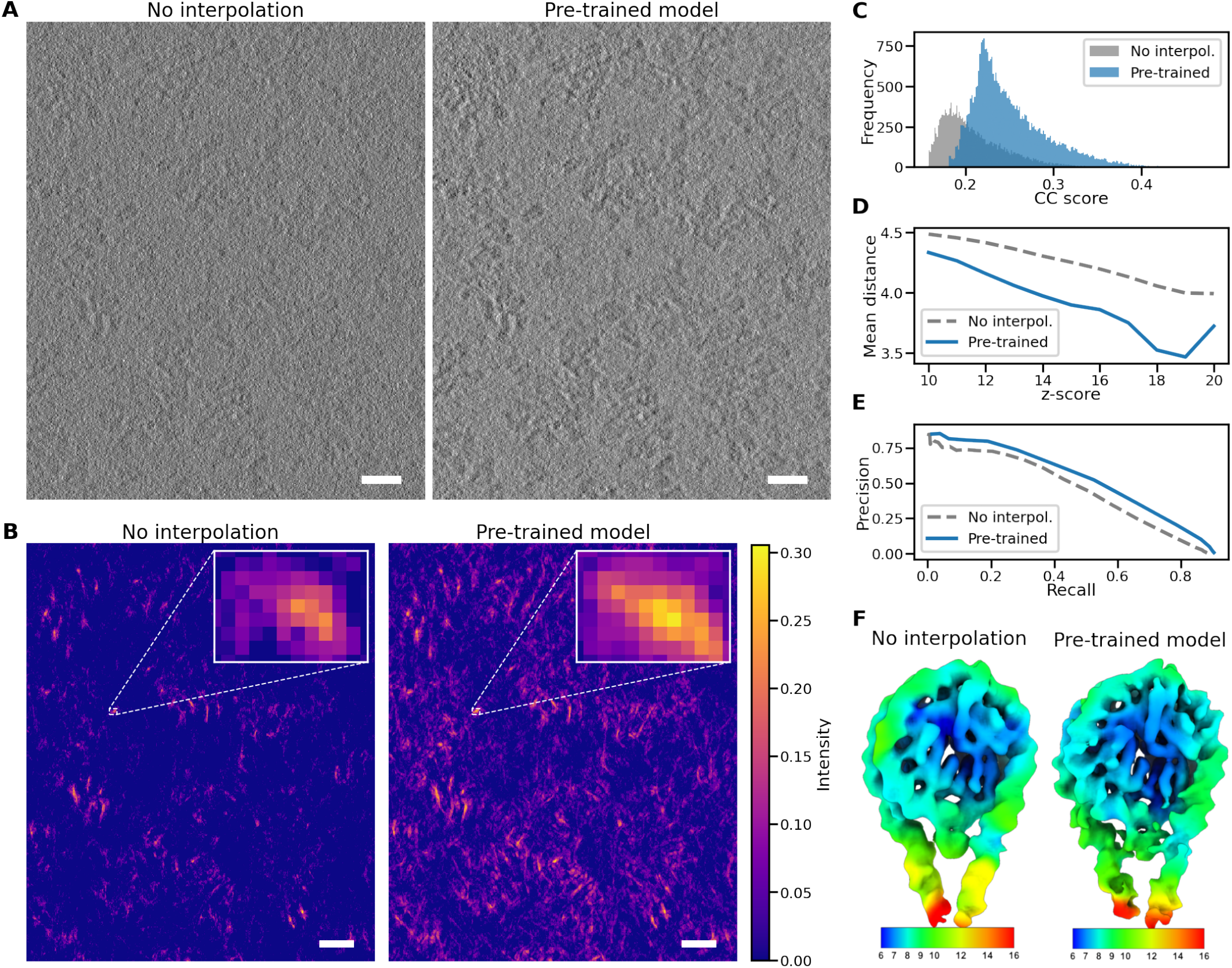
Enhanced particle picking and 3D structure determination. **(A)** Slice from the tomogram reconstructed without the interpolated tilt images (left) and with images interpolated using the pre-trained model (right). **(B)** Slices from TM results for the tomograms depicted in panel A showing improved contrast of the CC peaks for the interpolated data (right). **(C)** Histogram of CC scores from the selected peaks illustrating increase in absolute values as well as their frequency in case of the interpolated data. **(D)** Average distance of CC peaks to the refined positions found during STA showing improved precision in localization for the interpolated data. **(E)** Precision-recall curves with higher PR-AUC value for the interpolated data (0.4769) compared to non-interpolated data (0.4148). **(F)** 3D structure calculated based on non-interpolated (left) and interpolated positions colored by their local resolution (colorbar in Å). Although the structure based on the positions from interpolated data has similar resolution, it shows more enhanced structural details. The scale bar is 20 nm.

After running 3D TM in GAPSTOP^TM14^ on these tomograms, more distinct cross-correlation (CC) peaks with generally higher values were obtained, which makes them easier to identify (Fig. 4B,C). Since there was no GT for nucleosome positions, it was necessary to validate that the TM positions with the highest CC scores correspond to actual nucleosome locations in the tomograms. The authors^42^ validated the nucleosomes positions by manually creating a binary mask to separate the nucleus from the non-nucleosome-containing cytoplasm. Subsequently, only the nuclear peaks that were thresholded by the 99 percentile of the cytoplasmic peaks were considered. The baseline for both the non-interpolated and interpolated versions was established in this manner^42^.

Overall, in the 14 tomograms, we detected ∼18k nucleosome particle positions in the non-interpolated condition, while the interpolated variant identified ∼33k positions. We want to emphasize that we used interpolated data to detect positions, but the actual particles for STA were extracted from the tomograms without interpolation. Interpolated data could potentially contain additional high-resolution details that were not experimentally verified. Therefore, we use them to improve the reconstruction pipeline, but in order to determine the 3D structure, only experimentally acquired data are utilized.

The average distance between the matches in non-interpolated and interpolated peak positions confirmed that the latter are closer to the positions obtained after STA refinement and hence more precise in terms of their localization (Fig. 4D). We computed the precision-recall curves for both tested conditions (Fig. 4E), where we observe increase in PR-AUC value from 0.4148 for non-interpolated data to 0.4769 for interpolated data. A higher PR-AUC indicates that the model is better at correctly identifying positive instances without producing a large number of false positives.

When ∼33k nucleosome particle positions from the interpolated variant were used in the STA, it led to only marginal improvement in resolution (from 8.4 Å to 8.3 Å, see Supplementary Fig. 8). However, the map obtained from the interpolated-based particle list has more pronounced structural details as shown in Fig. 4F. The enhanced details are especially visible on the DNA linker arms. This use case shows great potential of interpolation for reliable localization of small features in crowded cellular environments (increase in founded positions by 87.33 % supported by 14.97 % improvement in PR-AUC).

### 2.5 Refined deep-learning particle picking

The aforementioned use cases primarily focus on downstream analyses using the template matching pipeline. Here, we demonstrate cryoTIGER application with DeePiCt^17^, an open-source deep-learning framework designed for supervised segmentation and macromolecular complex localization. DeePiCt models, trained on experimental cryo-ET data, are broadly applicable across species and datasets.

We used a Colab notebook provided by the authors^17^ to run the predictions of 3D convolutional neural network models on our tomograms. We used the model available in the Colab notebook that is trained for ribosome localization and performed predictions on five tomograms, both with and without interpolation. For the interpolation, we tested linear interpolation, the pre-trained model, and our model, which clearly exhibit a higher density of detected probability peaks compared to non-interpolated data (Fig. 5A), indicating improved ribosome localization.

**Figure 5.**
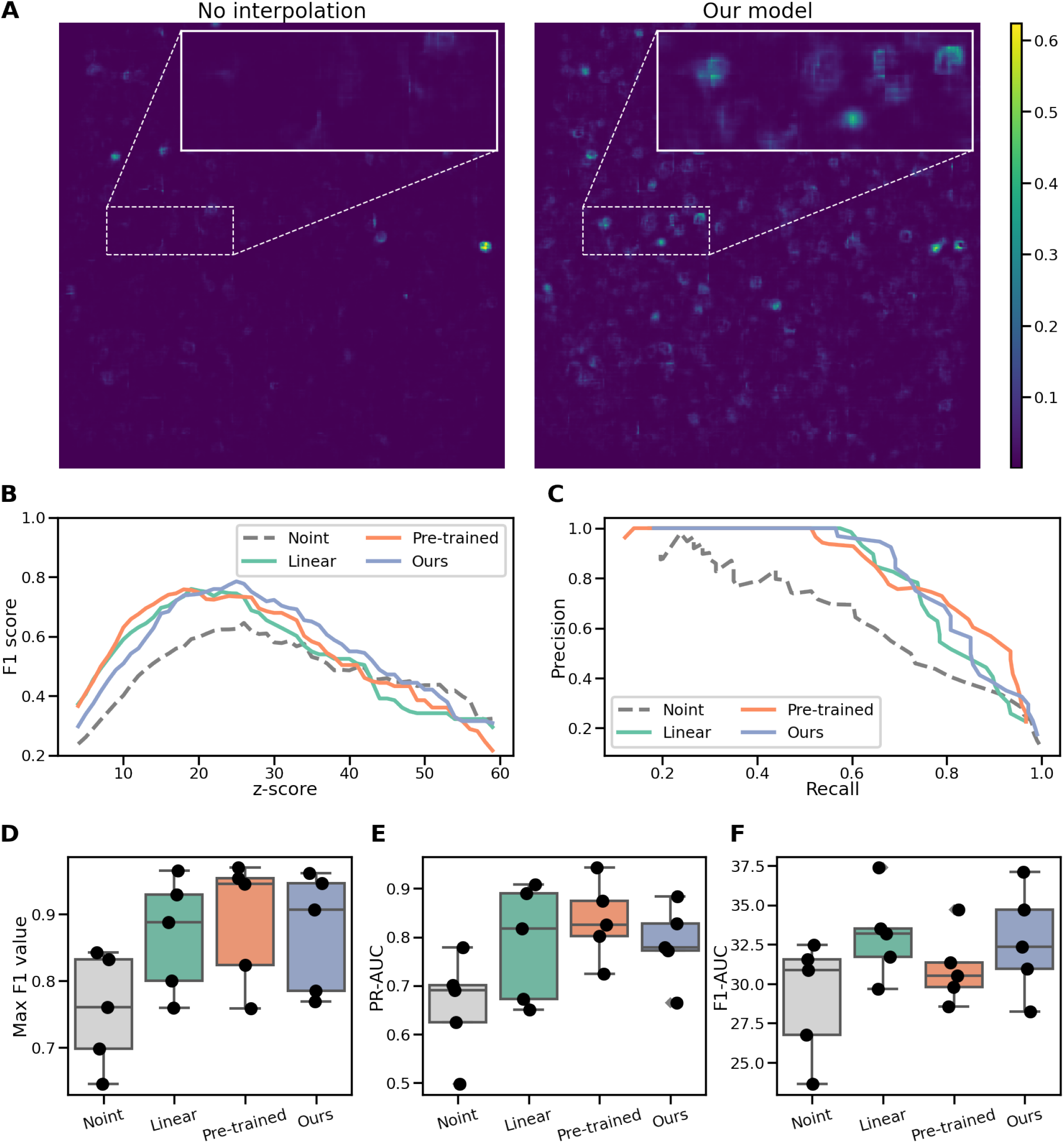
Picking ribosomes with DeePiCt. **(A)** 2D slice from the probability map generated by DeePiCt using non-interpolated data (left) and using interpolation with our model (right). The probability bar is identical for both maps. **(B)** F1 score for one representative tomogram. **(C)** Precision-recall curve comparison for one representative tomogram. **(D)** Maxima of F1 scores for five tested tomograms. **(E)** The area under the precision-recall curve for five tested tomograms. **(F)** The area under the F1-scores curve for five tested tomograms.

To validate founded positions, we compared them with a manually curated GT list from the Section 2.3 and compared the results using F1 scores, precision, and recall. The interpolated predictions consistently achieved higher values (Fig. 5B,C). To further substantiate these findings statistically, we used metrics such as the maximum F1 score, the area under the F1 curve (F1-AUC), and PR-AUC. All results, including those from linear interpolation and the cryoTIGER-based workflow, consistently show enhanced reconstruction localization properties when interpolated tilts are incorporated (Fig. 5D-E). This demonstrates that cryoTIGER provides improvements in particle localization that go beyond the TM approach.

### 2.6 Enhanced membrane segmentation

Lastly, we evaluated the impact of the interpolation on DL-based, fully automated membrane segmentation as implemented in MemBrain v2^43^. A core module of MemBrain v2 employs a 3D-UNet architecture optimized for cryo-ET membrane segmentation where the provided models were trained on diverse cryo-ET data, resulting in robust and widely-used software for membrane segmentation^44–46^.

To source publicly available annotated data in standardized formats, we utilized the cryoET Data Portal^47^. In the portal, we identified four tilt series (128_2, 129_2, 133, 141_3) from the dataset CZCDP-10004 containing all necessary reconstruction files as well as a hybrid segmentation mask to validate our method. The hybrid annotation method combines tomogram denoising, 3D-UNet-based membrane segmentation, and manual post-processing. We deliberately selected this hybrid approach to more accurately compare the performance of fully automated MemBrain v2 on non-interpolated and interpolated datasets.

We ran MemBrain v2 on the four tomograms reconstructed without interpolation, with the linear interpolation, and interpolation based on the pre-trained model as well as on our model. We assessed the quality of segmentation using both overlap-based and pixel-based metrics. The overlap-based metric, the Dice Similarity Coefficient, weighs the overlap twice as strongly as the non-overlap, while the Jaccard Index directly measures the proportion of overlap relative to the union of the ground truth and the segmentation (see Methods 4.4 for more details). The precision and recall where used to asses the segmentation on the pixel level. For all four metrics, we observed clear and consistent improvement in MemBrain v2 outputs on cryoTIGER-based workflow compared to linear interpolation or non-interpolated data (see Fig. 6). After interpolated frames from DL models were added, some false positive segmentation artefacts were removed (red color in panel B) and the automated segmentation becomes more closer to the GT hybrid annotation with less false negatives (green color in panel B). These results strongly demonstrate the potential of interpolation to enhance fully automated membrane segmentation.

**Figure 6.**
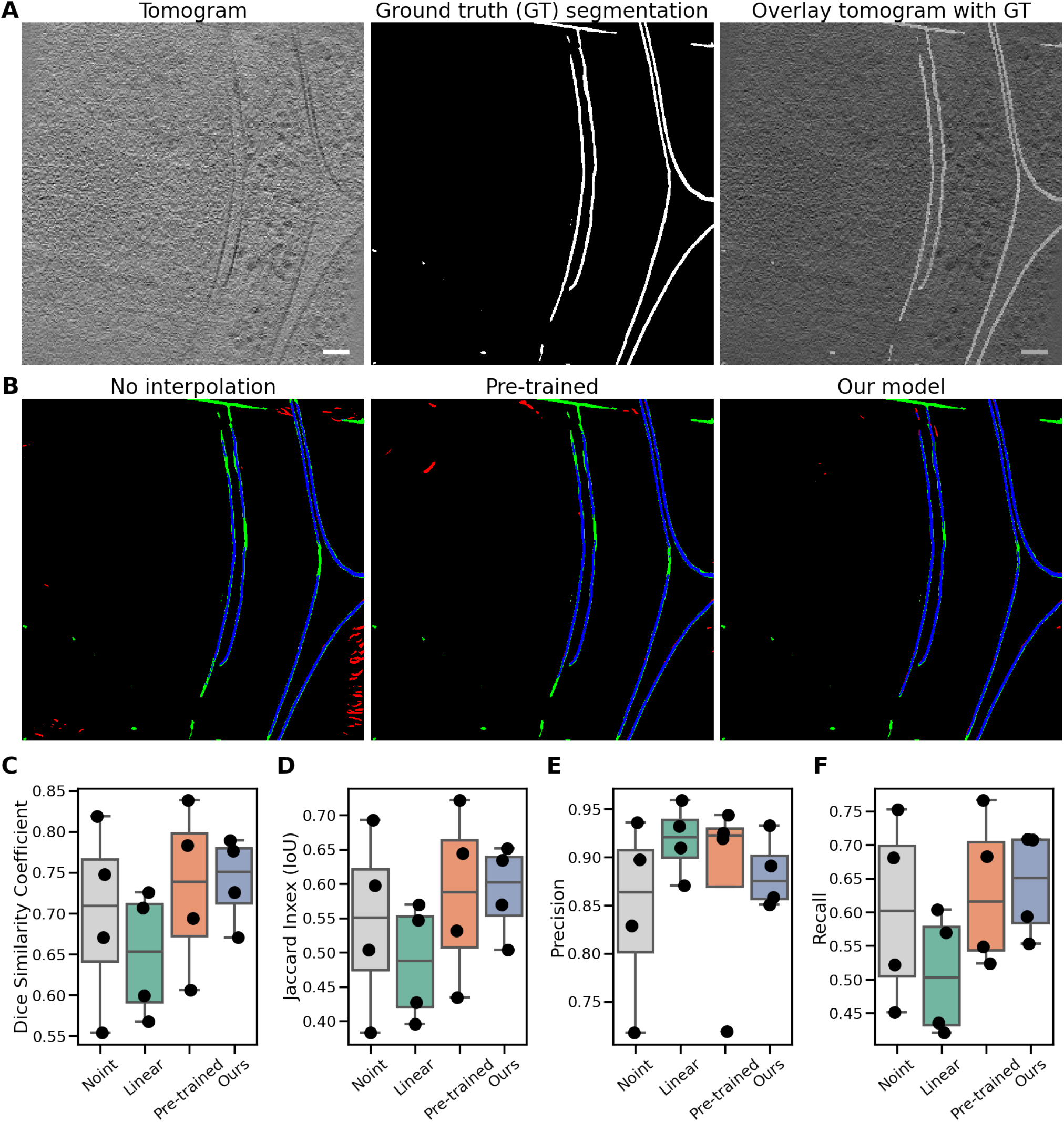
Enhanced membrane segmentation using interpolated tilt images. **(A)** A representative tomogram slice from cryoET Data Portal^47^ reconstructed using our pipeline (left). The corresponding slice showing the hybrid ground truth segmentation in the green channel (middle). The tomogram slice overlayed with the GT segmentation (right). **(B)** Comparison of non-interpolated version with interpolation using pre-trained model and our model trained on cryo-ET data. Green color represents areas in GT but not selected with MemBrain v2 on this tomogram (false negatives), red areas were selected by MemBrain v2 but are not in GT (false positives), blue areas were selected by both MemBrain v2 and GT (true positives). **(C)** Overlap-based statistics for Dice Similarity Coefficient for the four tested tomograms. The coefficient values ranged from 0 (no-overlap) to 1 (full overlap). **(D)** Overlap-based statistics for Jaccard Index for the four tested tomograms. The coefficient values ranged from 0 (no-overlap) to 1 (full overlap). **(E)** Pixel-based statistics for precision for the four tested tomograms. **(F)** Pixel-based statistics for recall for the four tested tomograms. Precision and recall should always be considered together, because when linear interpolation has higher precision and lower recall, it means that the model is good at identifying membrane when it makes a prediction, but it misses many membrane locations overall. Scale bar is 60 nm.

## 3 Discussion

To address the challenge of angular spacing in tomographic reconstructions, our study introduces a novel application of an existing frame interpolation framework, tailored to the cryo-ET pipeline. By interpolating images between acquired tilts, our approach effectively increases angular sampling, enhancing the signal content in the reconstructions. To ensure robustness, we developed a custom model through extensive training on diverse datasets, varying in both cellular content and acquisition setup. Validation experiments demonstrated that DL-based interpolation generates images that enhance tomogram reconstruction and outperforms in that sense the traditional linear interpolation.

The impact of the interpolated images on tomogram properties was comprehensively evaluated using template matching, DL-based particle localization with DeePiCt, and DL-based segmentation using MemBrain v2 across diverse datasets and targets. The results for DL-based interpolation consistently surpassed those obtained from the linear interpolation, with notable improvements observed in all cases. This was surprising given the overall good performance of the linear interpolation in 2D. One of the explanation could be that the DL-based approaches better preserve structural content (see Fig. 2F). While our model often excelled, there were instances where the pre-trained model performed better, underscoring the method’s inherent robustness and potential for improvement with targeted training.

The most notable advancements were achieved in challenging targets, such as the nucleosome dataset, where our method more than doubled the number of reliably localized particles in comparison to the non-interpolated data. Additionally, the automated membrane segmentation results showed greater agreement with manual annotations, highlighting the potential for reduced manual curation. This has far-reaching implications as accurate segmentation of membranes and the localization of associated proteins are essential for advancing cryo-ET studies that link membrane architecture and molecular organization to cellular function. Furthermore, if the automated segmentation is reliable enough, it could replace manual annotations of surfaces needed for surface-based particle localization of pleomorphic assemblies for subtomogram averaging.

The interpolation workflow may influence cryo-ET data acquisition parameters. It enables the use of larger tilt increments with increased electron dose per image without compromising the achievable content. This can improve the performance of downstream processing due to increased SNR. Moreover, for samples that are unusually sensitive to electron-dose, an adjusted tilt-acquisition-scheme can be combined with tilt interpolation, so data can still be acquired with a reasonable tilt range and sufficient electron-dose per image.

Although our study demonstrates the potential of interpolation approaches, it is not without limitations. While interpolated tilts can reduce small gaps in angular sampling, they do not resolve structural data loss associated with missing data across a larger angular space. Attempts to generate more than one interpolated image between two experimental tilt images using the current architecture occasionally resulted in artifacts, particularly when the interpolated tilt images deviated from realistic structural representations (see Supplementary Fig. 1). This highlights the need for careful optimization and evaluation of interpolation outputs to avoid introducing misleading features into reconstructions.

Future work should explore alternative neural network architectures beyond the FILM model to further optimize performance and address existing limitations. For instance, networks designed specifically for extrapolation may hold promise for mitigating the effects of the missing wedge. However, this poses a significant challenge, as it requires the development of approaches that can generalize well without over-fitting, especially in the absence of ground-truth data for validation. The integration of extrapolation networks or hybrid models capable of interpolating and extrapolating tilt series data could potentially open new ways for addressing this longstanding issue in cryo-ET.

In conclusion, our study underlines the importance of filling in the angular space between the tilt images and provides a novel computational solution for this problem. The presented interpolation approach has shown promising results, enhancing tomogram properties relevant for both particle and feature localization. To facilitate further research and community adoption, we provide our approach as an open framework cryoTIGER, complete with trained models, laying a solid foundation for the continued exploration of interpolation techniques. By addressing current limitations and pursuing innovative methods, we anticipate further advancements in the tilt-interpolation methods that will continue to enhance cryo-ET reconstructions and facilitate their analyses, thereby advancing structural biology studies.

## 4 Methods

### 4.1 Preprocessing

The input in our preprocessing pipeline is dose-filtered and aligned tilt series (for the detailed workflow from the raw tilt series to the aligned ones, we refer reader for example to the studies on 80S ribosome^41^ and NPC NR^40^). To accommodate memory constraints, the input tilt series are binned by a factor of 2.

Linear interpolation is performed by computing a pixel-wise average between each pair of adjacent tilt images. For completeness, we also considered the effect of tilting, where the pitch angle is determined as 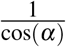. For tilt steps below 3 degrees, the pitch between two neighboring images is below 1, making linear interpolation between corresponding pixels valid. For tilt steps of 4 degrees, the pitch is approximately 1.2. In this case, one could consider computing linear interpolation between pixels with an offset of 1 or using cubic interpolation, which takes neighboring pixels into account. However, the results shown in the 3 suggest that linear interpolation without an offset was superior even for 4-degree tilt increments. This is most likely due to the downsampling of the data, which combines values from neighboring pixels, thereby diminishing the effects of the pitch. In our study, we only evaluated data with tilt increments below 4 degrees, so this pitch was not considered for linear interpolation.

In the deep learning-based interpolation process, the network requires input data with three color channels. Therefore, the grayscale data are normalized to the 0–255 range, copied into all three channels, and saved in the PNG format.

After executing the interpolation process, the output is generated in the RGB format. Converting this output into a grayscale image using the luminance channel involves combining the three color channels into a single intensity channel that represents the perceived brightness of each pixel. The formula for converting an RGB value to grayscale using the luminance channel is:

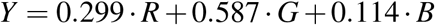

Finally, the image stack is reconstructed using the NumPy library and written out using cryoCAT^48^.

### 4.2 2D image comparison metrics

To assess the similarity between images we used PSNR, RMSE, and SSIM metrics. The PSNR is a measure used to assess the quality of a reconstructed image compared to the original image. It represents the ratio between the maximum achievable power of a signal and the power of noise that disrupts its accuracy. Given that many signals span a broad dynamic range, PSNR is typically measured on a logarithmic scale in decibels (dB). PNSR s given by the formula:

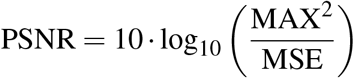

where MAX is the maximum possible pixel value of the image (e.g., 255 for 8-bit images) and MSE is the Mean Squared Error, calculated as:

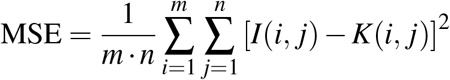

Here, *m* and *n* are the dimensions of the image, *I*(*i, j*) is the pixel value of the original image at position (*i, j*), and *K*(*i, j*) is the pixel value of the reconstructed image at position (*i, j*).

A higher PSNR value indicates better quality, with less distortion between the original and reconstructed images. PSNR is logarithmic, and a difference of 1 dB represents a multiplicative change in the MSE. Specifically, a difference of 1 dB indicates that the ratio of the peak signal power to the noise power has changed by approximately 26 %. This means the noise level in the image has either increased or decreased accordingly.

The RMSE is a metric used to quantify the difference between reconstructed values and actual values. It gives higher weight to larger errors, making it sensitive to outliers and it is calculated as the square root of the average of the squared differences:

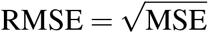

The RMSE is always non-negative, with lower values indicating smaller errors and better performance. RMSE directly measures the error, so a difference of 1 unit in RMSE means that, on average, the deviation between the processed and original image changes by 1 unit in the same scale as the data.

The SSIM is a metric used to assess the similarity between two images by focusing on structural information rather than pixel-wise differences. SSIM combines luminance, contrast, and structural comparisons into a single score ranging from −1 (completely dissimilar) to 1 (perfect similarity).

The formula for SSIM is:

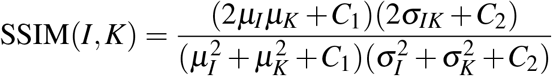

where *µ*_*I*_ is the mean intensity of image *I, µ*_*K*_ is the mean intensity of image 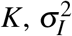 is the variance of image 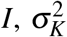 is the variance of image *K, σ*_*IK*_ is the covariance of images *I* and *K, C*_1_ = (*K*_1_ · *L*)^2^ is the stabilization constant for luminance, *C*_2_ = (*K*_2_ · *L*)^2^ is the stabilization constant for contrast, *L* is the dynamic range of pixel values (e.g., 255 for 8-bit images), and *K*_1_ and *K*_2_ are small constants (default values: *K*_1_ = 0.01, *K*_2_ = 0.03).

SSIM considers the luminance (comparing the mean intensity values), contrast (comparing the variances of the images), and structure (comparing the covariance of the images). A higher SSIM value indicates greater similarity between the two images. A difference of 0.01 in SSIM (e.g., from 0.85 to 0.86) indicates a 1 % change in structural similarity and larger differences (e.g., from 0.70 to 0.80) reflect a substantial improvement in similarity.

### 4.3 Metrics for evaluating peak selection

To assess the performance of particle selection, we computed the F1 score, precision, recall, and the PR-AUC between found positions and the GT lists.

In order to evaluate TM results with and without interpolation, we extracted peaks with different thresholds and compared them against baseline positions. When evaluating the match between a list of found peak positions and the baseline positions, we introduced a tolerance distance to account for imprecisions arising from particle list conversions. A peak was considered a match if it was found within this tolerance distance from the baseline position.

Precision measures the proportion of correctly predicted positive samples among all positive predictions. It is defined as:

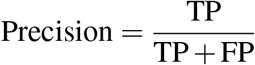

where TP are True Positives (correctly predicted positive samples) and FP are False Positives (incorrectly predicted positive samples).

Recall (also known as Sensitivity) measures the proportion of correctly predicted positive samples among all actual positive samples. It is defined as:

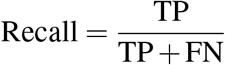

where FN are False Negatives (actual positives incorrectly predicted as negative).

The F1 Score is the harmonic mean of precision and recall, providing a single measure that balances the two metrics. It is defined as:

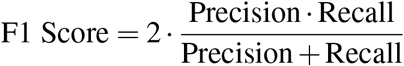

The Precision-Recall AUC is the area under the precision-recall curve, which plots precision (*y*-axis) against recall (*x*-axis) for different thresholds. It is computed as:

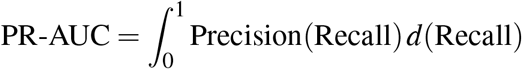

Here, the integral is calculated over the range of recall values from 0 to 1.

### 4.4 Contrast-based metrics for visual quality

Standard Deviation Contrast is a measure used to quantify the contrast of an image by evaluating the variation in pixel intensities. It reflects how much the pixel values deviate from the mean intensity. A higher standard deviation indicates greater contrast, while a lower value suggests lower contrast.

The standard deviation contrast is calculated as the standard deviation of the pixel intensities, given by:

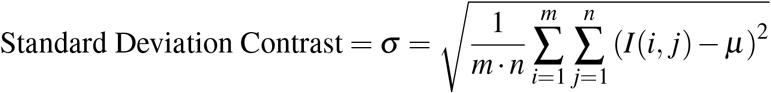

where *I*(*i, j*) is the pixel intensity at position (*i, j*) and *µ* is the mean intensity of the image, defined as:

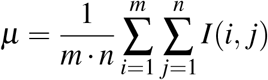

where *m* and *n* are the height and width of the image, respectively.

A higher standard deviation of pixel intensities indicates higher contrast in the image, whereas a lower standard deviation suggests the image has less contrast.

Gradient-Based Contrast is a measure used to quantify the contrast of an image based on the spatial variations in intensity between neighboring pixels. This metric highlights areas with sharp intensity changes, such as edges, and is useful for evaluating the sharpness and detail of an image.

The Gradient-Based Contrast is calculated by summing the gradient magnitudes of the image in the horizontal (*x*) and vertical (*y*) directions. The formula is given by:

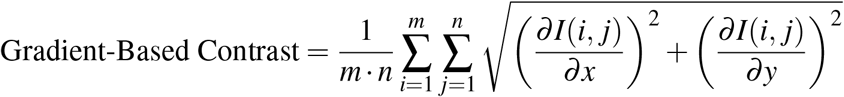

where 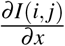 is the partial derivative of the image with respect to the horizontal direction (*x*), 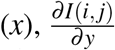 is the partial derivative of the image with respect to the vertical direction (*y*), and *m, n* are the height and width of the image, respectively.

The partial derivatives are often approximated using convolution filters, such as the Sobel filter^29^, to calculate the gradient in each direction. A higher gradient contrast value indicates a higher degree of contrast in the image, with sharper transitions between pixel intensities.

### 4.5 Experimental data

#### 4.5.1 Parameter setups for TM

In our TM configuration, we used NPC NR template (EMD-51628)^40^ with the low-pass filter of 23 pixels (corresponding to 30 Å) and angular sampling of 10 degrees on 4× binned data. For detecting ribosomes, we used 80S ribosome template (EMD-15807)^41^ with the low-pass filter of 30 pixels (corresponding to 23 Å) and angular sampling of 10 degrees on 4× binned data.

#### 4.5.1 Nucleosome template matching and subtomogram averaging

The nucleosome structures shown in Fig. 4 were obtained from 14 tomograms of resting T cells. The dataset is the main focus of a separate study^42^ and we therefore refer a reader to read the original work for details on the data acquisition parameters and processing. Here we briefly summarize information relevant for our study. The tilt series were acquired with tilt step of 2 degrees and angular range ±60 degrees resulting in 61 images and total electron dose of 135 e^*−*^ per Å. The pixel size of unbinned data was 1.971 Å.

To compare the performance of the cryoTIGER workflow for TM of nucleosomes, we used GAPSTOP^TM14^ on novaCTF^37^ reconstructed tomograms that contained either non-interpolated data or data interpolated with the pre-trained model. The TM was performed on data downsampled by factor of 2, i.e. pixel size of 3.942 Å.

The nucleosome template was the same for both cases, a lower resolution *in situ* nucleosome structure generated from the aforementioned 14 tomograms^42^. The peaks were extracted with the same thresholding approach^42^ and further cleaned by excluding clashing particles with a nucleosome shape mask around each peak in cryoCAT^48^.

The 17,916 (without interpolation) and 33,560 particles (with cryoTIGER workflow) determined through template matching were extracted as unbinned subtomograms in Warp^49^ (for both cases from non-interpolated data) and subjected to subtomogram averaging and alignment in Relion 3.1^50^. The particle set was then imported into M^51^ to perform multi-particle refinement of the tilt-series and the final structure. This resulted in a chromatosome structure (containing the core nucleosome with H1 and DNA linkers) resolved to 8.4 Å (with no interpolation) and 8.3 Å (with cryoTIGER workflow) where the latter structure contained more details (Fig. 4G,H).

Note that the chromatosome structure based on the particle list obtained from the interpolated data was further refined and classified, reaching the local resolution of 6.4 Å (7.3 Å overall)^42^. Corresponding Fourier shell correlation (FSC) curves are in Supplementary Fig. 8. This procedure was not reproduced with the particle list based on non-interpolated data due to extensive computational and time costs.

### 4.6 Time complexity and memory limitations

For input tilt series consisting of 61 tilts with the resolution of 2048 ×2048, the interpolation network runs for approx. five minutes on a machine with AMD EPYC 7543P 32-Core Processor and two NVIDIA A100 80 GB GPU cards. In terms of memory limitations, we could not fit the full unbinned data into memory. Therefore, we ran all DL interpolation tests on data binned by a factor of 2, which corresponds to the aforementioned resolution.

## Data availability

The primary data from the ribosome and NPC NR evaluation are publicly available on EMPIAR (EMPIAR-11899, EMPIAR-12454, respectively). The tomograms used for DL-based segmentation are available at CZI data portal - the dataset ID is CZCDP-10004 and tilt series 128_2, 129_2, 133, 141_3 were used for the evaluation.

## Code availability

The framework cryoTIGER is publicly available on GitHub https://github.com/turonova/cryoTIGER/ in form of Jupyter notebook (under GPL-3.0 license). All trained models as well as minimal example on how to run the interpolation on tilt series are provided there as well.

## Acknowledgements

We thank Desislava Glushkova and Huaipeng Xing from Max Planck Institute of Biophysics for providing data for the training of the models. We thank Stefanie Böhm for critically reading of the manuscript and helpful discussion. We thank Thomas Hoffman from EMBL and the Max Planck Computing and Data Facility for support with scientific computing. All data in this manuscript were acquired at the Central Electron Microscopy Facility at Max Planck Institute of Biophysics.

## Author information

Conceptualization: T.M, and B.T. Investigation: T.M, and B.T. Methodology: T.M. and B.T. Data curation: T.M, J.P.K, M.W.T., and B.T. Validation: T.M. and B.T. Formal analysis: T.M., J.P.K, J.L., and B.T. Software: T.M. and B.T. Project administration: T.M. and B.T. Visualization: T.M., J.P.K, S.C.-L. and

B.T. Supervision and funding acquisition: G.H., M.B., and B.T. Writing—original draft: T.M. and B.T. Writing—review & editing: T.M., J.P.K., M.W.T., S.C.-L., J.L., G.H., M.B., and B.T.

