## Supplementary Information for "cryoTIGER: Deep-Learning Based Tilt Interpolation Generator for Enhanced Reconstruction in Cryo Electron Tomography"

| # of TS | Tilt increment | # of triplets | # of iterations | Organism (image proc. method) |
|---|---|---|---|---|
| **375** | **1+2+3** | **317,312** | **up to 3mil** | **All from Table 1** |
| 285 | 1+2+3 | 14,987 | up to 2mil | *D. discoideum* (bin4 data) |
| 285 | 1+2+3 | 239,791 | up to 2mil | *D. discoideum* (patches from bin4 data) |
| 306 | 3 | 166,175 | up to 5mil | *D. discoideum* (patches from bin4 data) |
| 52 | 1 | 5,318 | up to 5mil | *D. discoideum* (bin4 data) |
| 119 | 2 | 6,665 | up to 5mil | *D. discoideum* (bin4 data) |
| 159 | 3 | 5,678 | up to 5mil | *D. discoideum* (bin4 data) |
| 171 | 1+2 | 12,343 | up to 5mil | *D. discoideum* (bin4 data) |
| 32 | 2 | 28,976 | up to 1mil | Human T cells (bin4 data) |
| 32 | 2 | 66,996 | up to 1mil | Human T cells (patches from bin4 data) |
| 32 | 2 | 66,996 | up to 3mil | Human T cells (deconvolve) |
| 32 | 2 | 66,996 | up to 3mil | Human T cells (topaz) |

**Supplementary Table 1.** Summary of tested models and configurations. Models with fewer number of iterations than is specified were also stored during the training. Training was done either directly on bin4 data or on 256×256 patches as described in the main text.

### FFT artefacts during training of our model

When training our model, we examined the Fourier transform (FFT) of the generated tilt images to visualize the presence of signals across different wavelengths. As a reference, we used a microscopic tilt image and its FFT (Sup. Fig. 4a), where no visible artifacts were observed. We then attempted to train the model using *D. discoideum* images with different data binning (Sup. Fig. 4b,c), a decreased learning rate (Sup. Fig. 4d), and transfer learning with weights from a pre-trained model (Sup. Fig. 4e). Additionally, we experimented with data from Human T cells (Sup. Fig. 4f) and extracted 256×256 patches from larger inputs (Sup. Fig. 4g,h). Each of these tests introduced specific artifacts in the frequency domain.

To better understand the causes of these distortions, we analyzed the original RGB Vimeo-90k dataset[38] (Sup. Fig. 5a) and applied a series of modifications to observe their effects in the frequency domain. Converting the RGB inputs to grayscale (Sup. Fig. 5b) and adding artificial CTF deformation (Sup. Fig. 5c) produced no visible differences. However, introducing Gaussian noise to the Vimeo-90k dataset (Sup. Fig. 5d) resulted in FFT artifacts similar to those observed in our cryo-ET data. This led us to conclude that the primary factor causing artifacts in our cryo-ET data is the presence of noise, likely in combination with other factors.

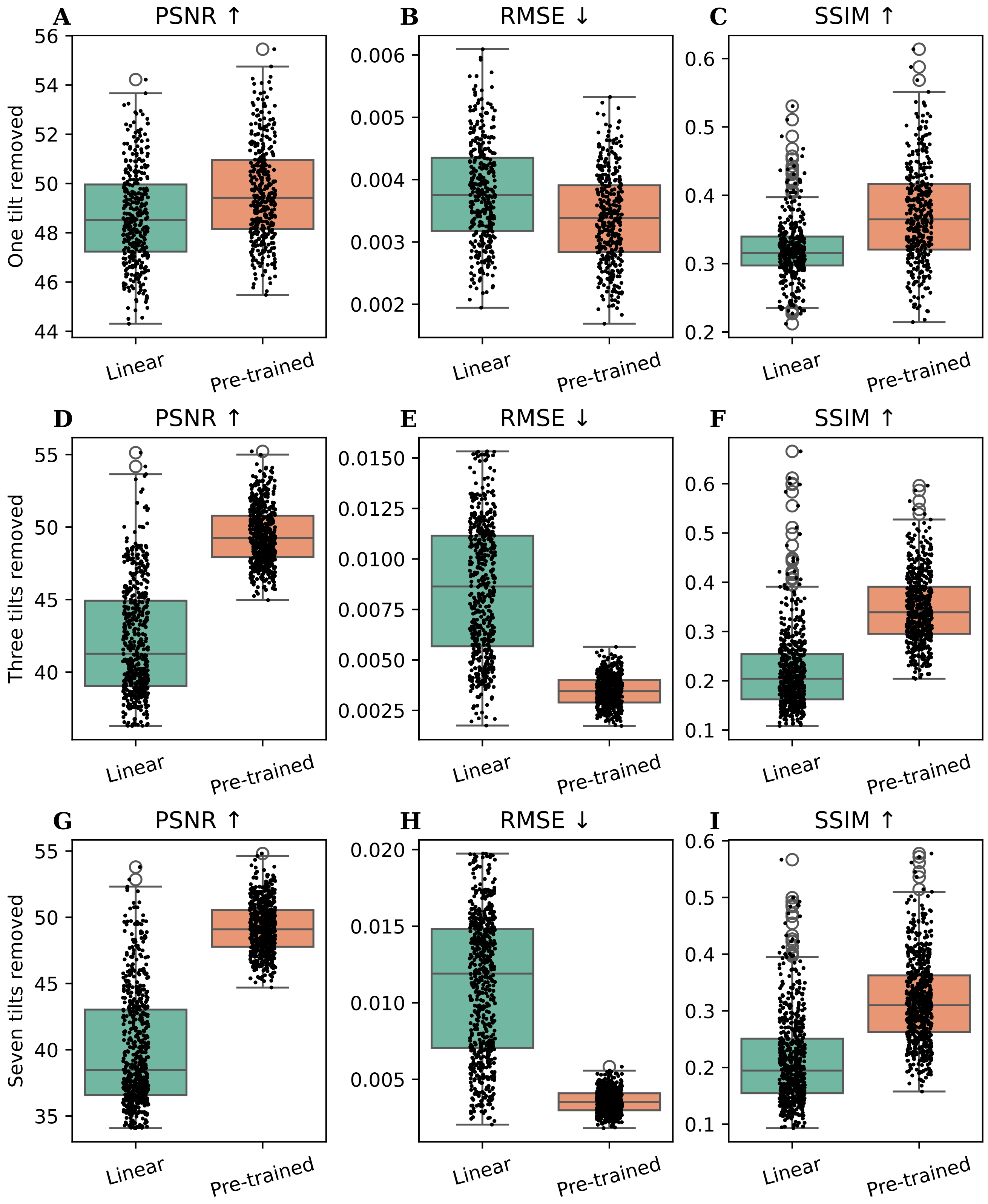


**Supplementary Figure 1.** Comparison of different interpolation methods when real tilts are removed. The test was done on same data as Fig. 2. Each boxplot contains data from seven tilt series binned by a factor of 8.

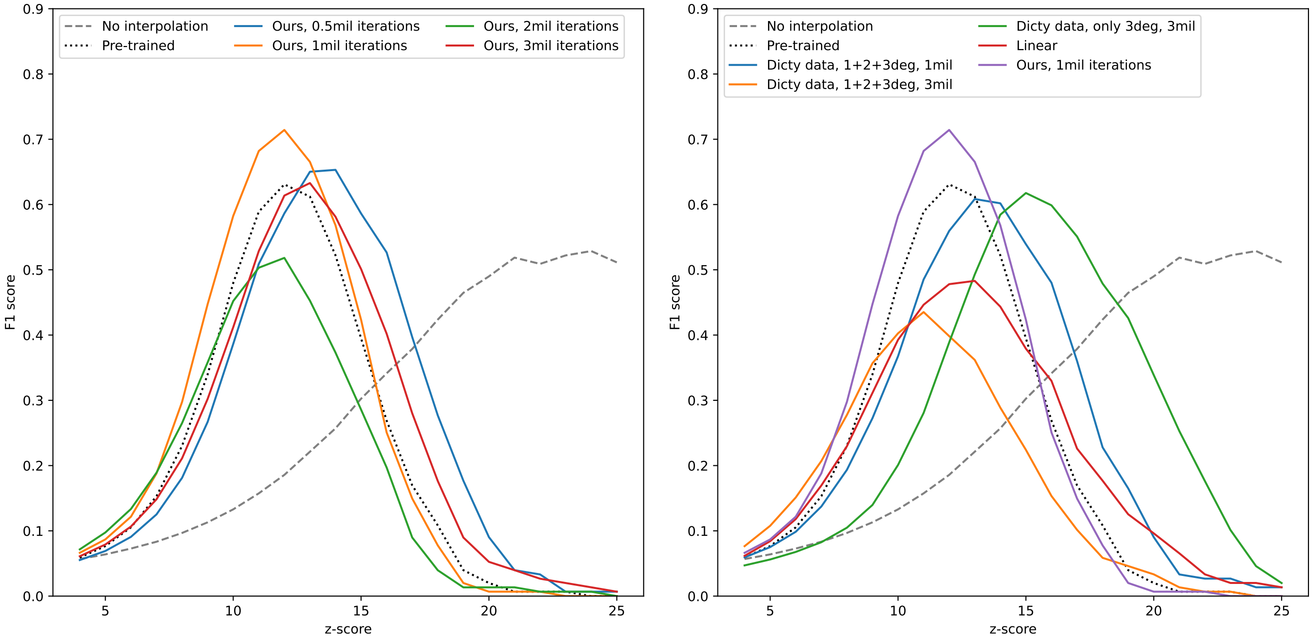


**(a)** The model training with different number of iterations. **(b)** Comparison of *our model* with other selected trained models.

**Supplementary Figure 2.** Particle picking results for 80S ribosome on different trained models.

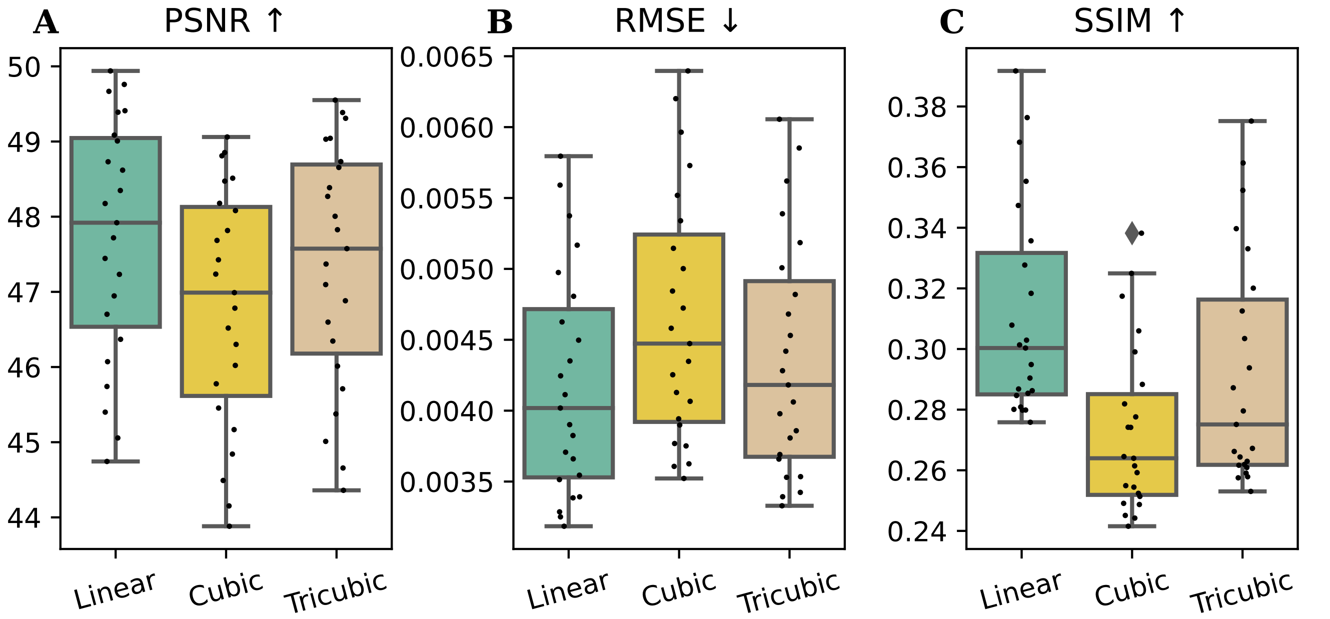


**Supplementary Figure 3.** Comparison between linear, cubic, and tricubic interpolation showing the best results for linear interpolation in all three metrics.

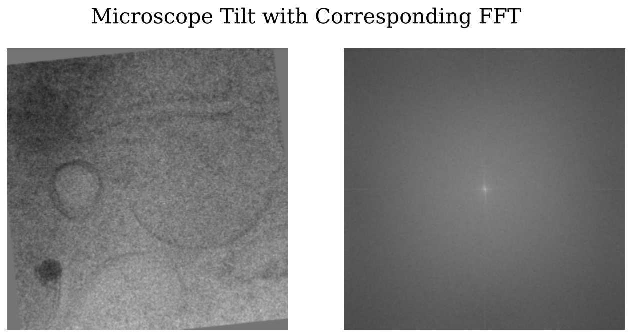


**(a)** Reference microscopic tilt image.

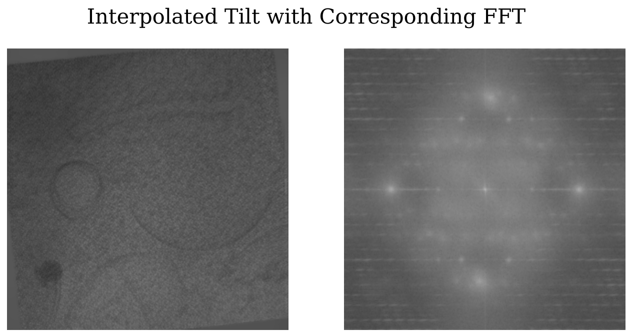


**(b)** Training on *D. discoideum*, bin4 input.

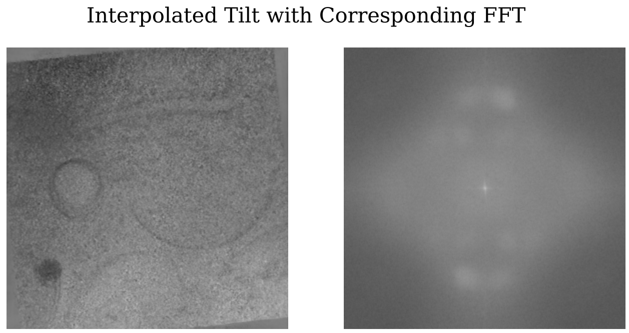


**(c)** Training on *D. discoideum*, bin16 input.

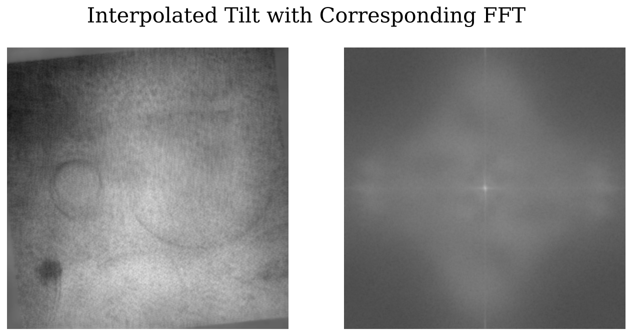


**(d)** Training on *D. discoideum*, decreased learning rate.

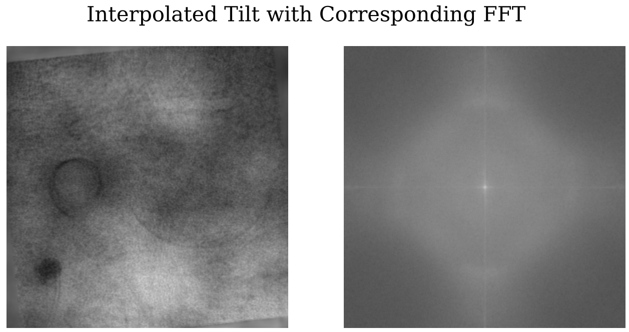


**(e)** Training on *D. discoideum*, transfer learning.

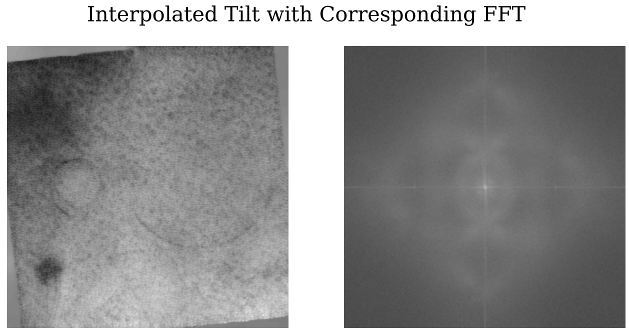


**(f)** Training on Human T cells, bin4 input.

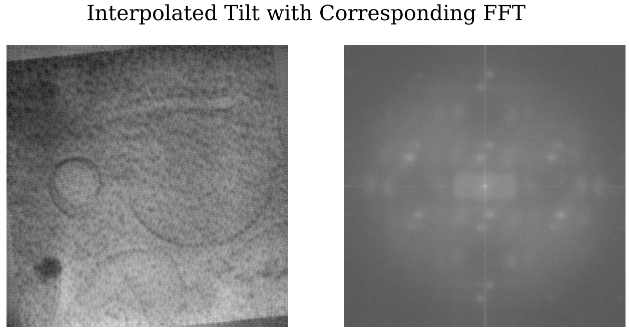


**(g)** Training on Human T cells, 256×256 patches.

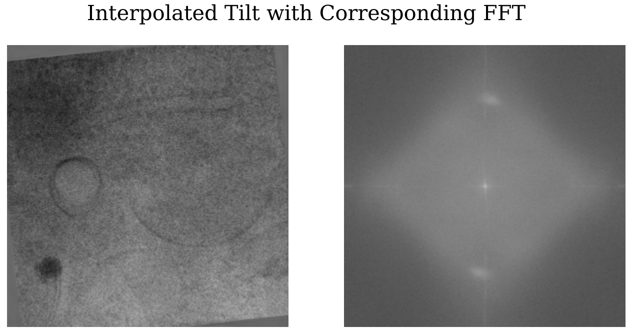


**(h)** Training on *D. discoideum*, 256×256 patches.

**Supplementary Figure 4.** The collection of FFT artifacts, when trained on cryo-ET data.

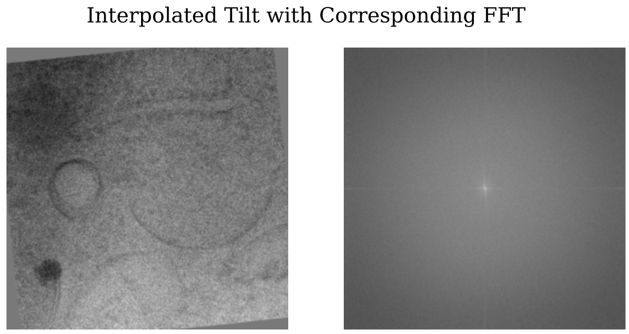


**(a)** Original RGB input in training.

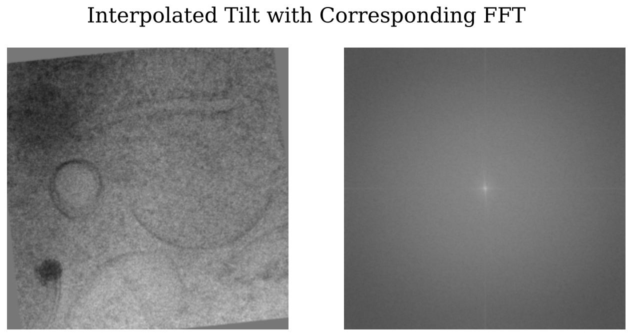


**(b)** Grayscale input in training.

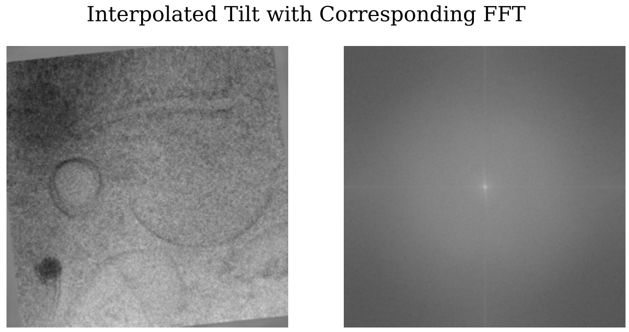


**(c)** Original RGB input plus CTF in training.

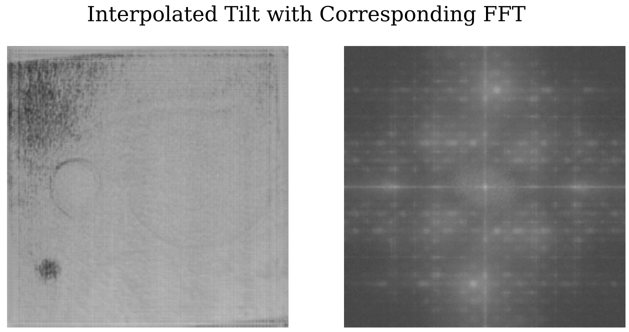


**(d)** Grayscale input plus Gaussian noise in training.

**Supplementary Figure 5.** The collection of FFT artifacts, when trained Vimeo-90k[38] dataset. Artifacts in training are result of multiple factors including noise.

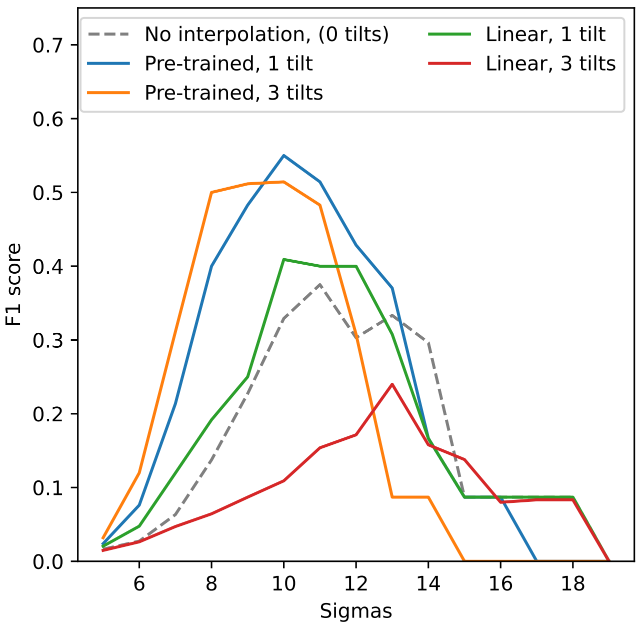


**(a)** NPC NR particle picking.

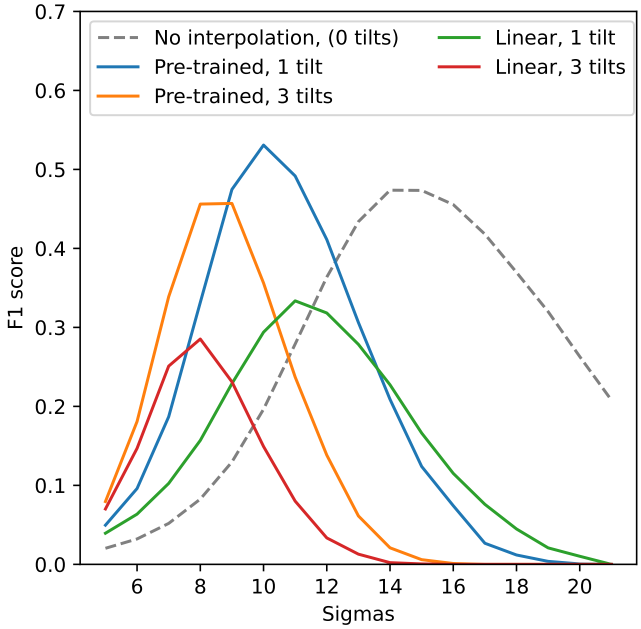


**(b)** Nucleosome particle picking.

**Supplementary Figure 6.** Comparison of adding one versus three interpolated tilts for different targets. For the nucleosome dataset, we compared the extracted list with the baseline, the formation of which is described in the main text. In both cases, using more interpolated tilts resulted in worse particle-picking performance.

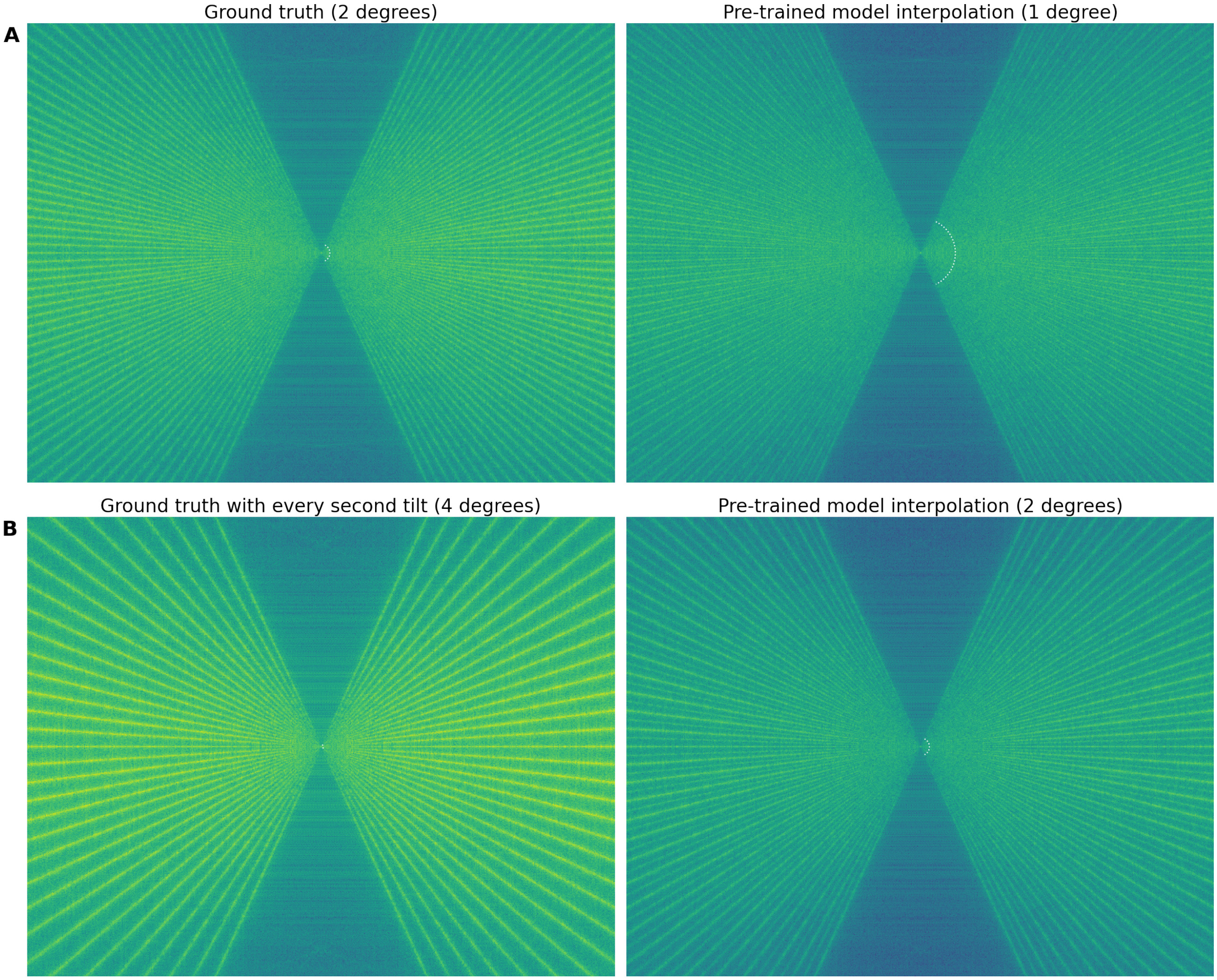


**Supplementary Figure 7.** Effects of the interpolation on angular sampling in reconstructed tomogram. The white dotted sections marks the low-resolution area where the angular sampling is still complete. The radii in Fourier pixels were computed using Crowther criterion formula. (A) Power spectrum of XZ tomogram slice of a reconstructed tomogram using the experimental tilt series with 61 images and 2 degrees tilt step. The complete angular sampling ends at radius 15.25 (in Fourier pixels). (B) Power spectrum of the same tomogram slice as in (A) but with additional tilts created by DL-based interpolation using the pre-trained model. The tilt series contains 121 images, tilt step is 1 degree, the complete angular sampling ends at 60.5 Fourier pixels. (A) Power spectrum of the same tomogram slice as in (A) with every second tilt removed. Tilt series contains 31 images, tilt step is 4 degrees, the complete angular sampling ends at 3.875 Fourier pixels. (D) (A) Power spectrum of the same tomogram slice as in (A) with every second tilt removed and then restored using DL-based interpolation with the pre-trained model. The tilt series contains 61 images, tilt step is 2 degree, the complete angular sampling ends at 15.25 Fourier pixels.

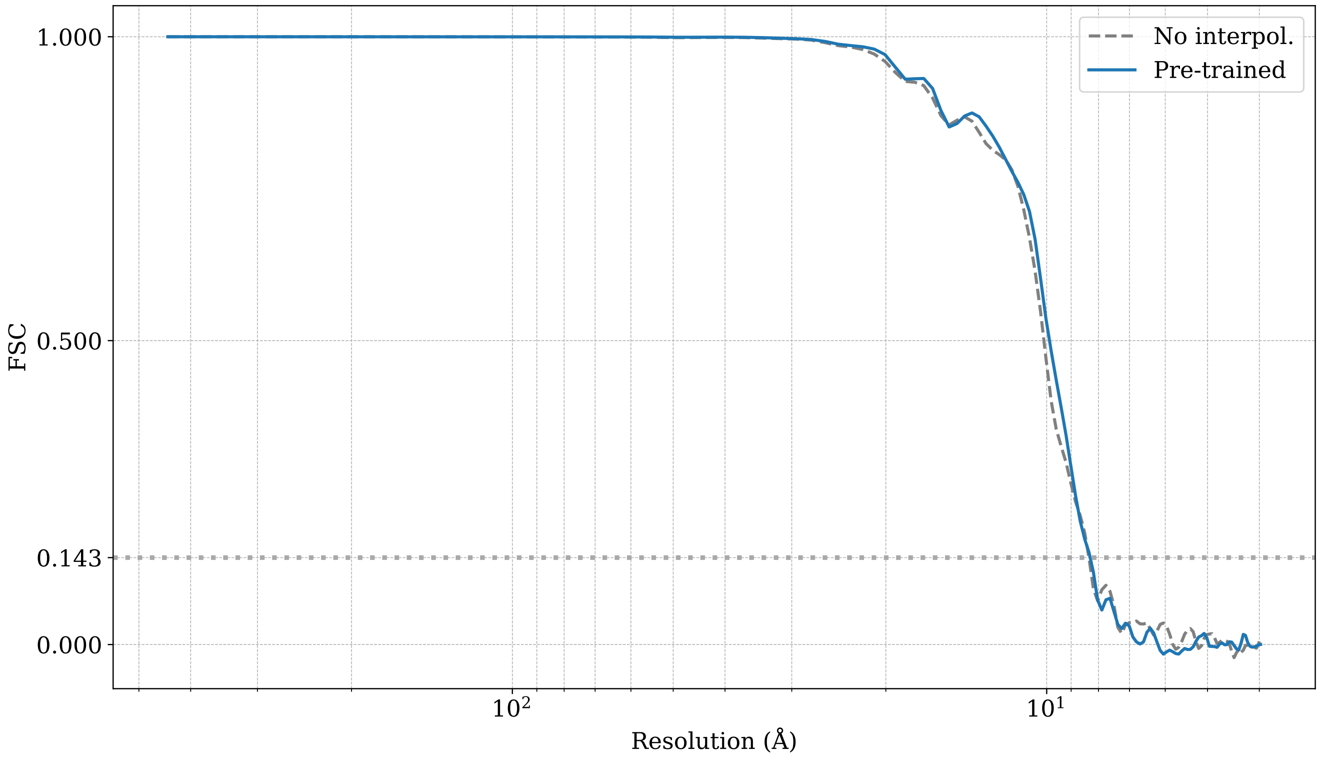


**Supplementary Figure 8.** Comparison of Fourier shell correlation (FSC) curves for a nucleosome structure determined using positions obtained from interpolated and non-interpolated data.
